# Estimating physical conditions supporting gradients of ATP concentration in the eukaryotic cell

**DOI:** 10.1101/2025.02.20.639229

**Authors:** Rajneesh Kumar, Iain G. Johnston

**Affiliations:** Department of Mathematics, University of Bergen, Bergen, Norway; Computational Biology Unit, University of Bergen, Bergen, Norway

## Abstract

The ATP molecule serves as an energy currency in eukaryotes (and all life), providing the energy needed for many essential cellular processes. But the extent to which substantial spatial differences exist in ATP concentration in the cell remains incompletely known. It is variously argued that ATP diffuses too quickly for large gradients to be established, or that the high rates of ATP production and use (sources and sinks) can support large gradients even with rapid diffusion – and microscopic models and detailed experiments in different specific cases support both pictures. Here we attempt a mesoscopic investigation, using reaction-diffusion modelling in a simple biophysical picture of the cell to attempt to ask, generally, which conditions cause substantial ATP gradients to emerge within eukaryotic cells. If ATP sources (like mitochondria) or sinks (like the nucleus) are spatially clustered, large fold changes in concentration can exist across the cell; if sources and sinks are more spread then rapid diffusion indeed prevents large gradients being established. This dependence holds in model cells of different sizes, suggesting its generality across cell types. Our theoretical work will complement developing intracellular approaches exploring ATP concentration inside eukaryotic cells.

## Introduction

Adenosine triphosphate (ATP) serves as a source of energy for many biochemical reactions throughout the cell. The concentration of ATP (and the ATP:ADP ratio, influencing the free energy available from ATP hydrolysis) determines the energy availability for a given reaction. ATP concentration in animal cells, for example, is often in the range 1-10mM (Greiner & Glonek, 2021). Cellular ATP concentrations vary through tissues and under different conditions (De Col et al., 2017). But how much does local ATP concentration vary within a cell? What influences whether substantial spatial gradients of ATP exist across cellular regions – and hence whether energy availability is in a sense heterogeneous throughout the cell?

As a small molecule (consisting of 47 atoms), ATP diffuses rapidly through the cytoplasm (of the order of several hundred μm^2^ s^-1^, sufficient to cross a 10μm cell in around 0.2s). But this does not necessarily mean that the profile of ATP concentration will equilibrate to a spatially uniform level. If localised sources and/or sinks of ATP produce and/or consume ATP at a sufficient rate, arbitrarily large concentration gradients can still exist.

Localised sources of ATP do exist in the eukaryotic cell. Mitochondria are bioenergetic organelles found in the majority of eukaryotic cell types (Roger et al., 2017) (Fig. 1A). Among many other processes (Picard & Shirihai, 2022), they catalyse the production of ATP from ADP and inorganic phosphate, through the process of oxidative phosphorylation (OXPHOS). OXPHOS is localized to mitochondria and often generates ATP at a greater rate than other cellular processes (including glycolysis), making mitochondria important local sources of ATP (though not under all conditions – see Discussion). Mitochondria are highly dynamic within many eukaryotic cells, moved by motor proteins on the cytoskeleton which themselves require ATP. An often-discussed question is the extent to which the speed of mitochondrial motion is directly controlled or a passive result of mitochondrial output providing this ATP supply.

**Figure 1.**
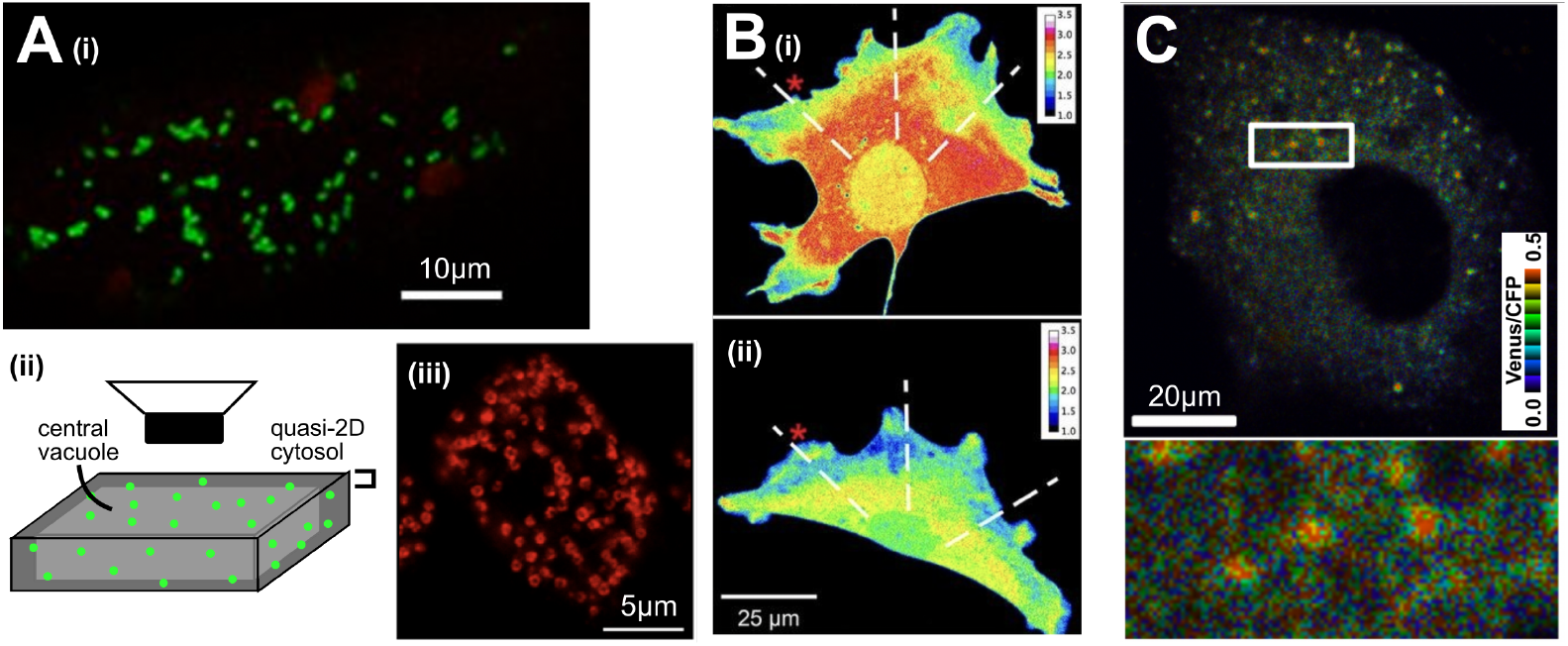
Example of cell geometries and ATP-gradient-related experimental results. **(A)** Examples of eukaryotic cell structures with punctate mitochondria. (i) An example plant cell from the hypocotyl of an *Arabidopsis thaliana* (plant) seedling, with discrete, punctate mitochondria visualized using green fluorescent protein (Chustecki et al., 2022; Logan & Leaver, 2000) (red objects are chloroplasts). (ii) A schematic structure of plant hypocotyl cells, where a large central vacuole makes the cytosol almost 2D (Lee Erickson et al., 2023); the image from (i) is imaged using a microscope with focal plane aligning with this section. (iii) A *Dictyostelium discoideum* (amoeba) cell with mitochondria visualized with anti-mitoporin from (Rai et al., 2011). **(B)** A mouse embryonic fibroblast (MEF) cell with ATP:ADP ratio visualized (heatmap) using the PercevalHR probe (Tantama et al., 2013). (i) shows a vertical view as in (A), (ii) shows a vertical section through the same cell, with smaller dimension and more uniform concentration profile. **(C)** A Huh-7 (immortalized epithelial) cell using the ATeam sensor (Imamura et al., 2009) to report ATP levels (heatmap). **Licensing:** (A) (iii) is taken from (Rai et al., 2011) under a CC-BY-NC license and has been edited to add annotations. (B) is taken from (Schuler et al., 2017) under a CC-BY-NC-SA license and has been edited to replace annotations; this subpanel of this figure is therefore also subject to a CC-BY-NC-SA license. (C) is taken from (Ando et al., 2012) under a CC-BY license and has been edited to move the colour scale and add annotation to the length scale bar.

Preserving spacing between mitochondria is hypothesized to be a priority in many cellular cases (Chustecki et al., 2021, Chustecki et al., 2022), particularly in the context of the cellular society (Chustecki & Johnston, 2024; Wang & Mukherji, 2022). Even spacing is valuable for uniform distribution of metabolites and signals through the cell and faithful partitioning of mitochondria at cell divisions (Aryaman, Bowles, et al., 2019; Edwards et al., 2021; Glastad & Johnston, 2023; Hoitzing et al., 2015; Jajoo et al., 2016; Moore et al., 2021).

ATP concentration features in a large collection of biomathematical models for metabolism and other cellular processes. Such models may use a well-mixed approximation, where concentrations do not vary with spatial coordinates (though may vary with time) and are in a sense averaged across the cell. Some examples include the modelling classes in (Beard & Qian, 2008; Forrest et al., 2023; Kerr et al., 2019; Kumar & Johnston, 2024), and indeed most “whole-cell models” (Goldberg et al., 2018). Other models explicitly represent the concentration of ATP (and associated metabolites) as a function of position within a specific cell type, often with a reaction-diffusion picture (Ghosh et al., 2018; Hatano et al., 2011, 2015; Vendelin et al., 2000). One recent example is the demonstration that spacing between mitochondria can emerge as a consequence of their influence on and response to the (heterogeneous) ATP profile of the cell in axons (Kajita et al., 2024). Another model, focusing on mitochondrial arrangement in cardiac cells, suggests that ATP concentration variability across the cell is limited in this system – under 10μM against an average concentration around 10mM – despite an uneven mitochondrial distribution (Ghosh et al., 2018). Similar reaction-diffusion approaches have been used to explore the cellular profiles of other related chemical species including oxygen (Sedlack et al., 2022) and calcium (Colman et al., 2022), and indeed of ATP within the mitochondrion (Garcia et al., 2019). Bridging these targeted approaches and the whole-cell modelling paradigm, spatial algorithms have been recently developed that consider the spatial behaviour of ATP (and other metabolites) with specific, detailed cellular structure (Francis et al., 2024).

Experimental characterization of the within-cell concentration profiles of ATP (and associated metabolites) is typically through fluorescence microscopy (recently reviewed in (San Martín et al., 2022)). Many beautiful approaches have revealed bulk or compartment-specific ATP levels (for example, comparing cytosolic to mitochondrial concentrations); we focus here on those that directly report *spatial variation* in ATP concentration in the cytosol. Groundbreaking work has revealed how mitochondrial position in mouse embryonic fibroblasts directly influence the spatial structure of ATP:ADP ratio within the cell, with the ratio highest in regions of dense mitochondria and with around 3-fold variation through the cell (Schuler et al., 2017) (Fig. 1B). In agreement, the ATP:ADP ratio in the quasi-1D system of mouse axons decreases with distance from mitochondria (Matsumoto et al., 2022), which modelling suggests can lead in term to emergent uniform spacing of mitochondria along the axon (Kajita et al., 2024). Semi-quantitative visualization of ATP levels in HEK293 (immortalized kidney) cells also show moderate within-cell ATP differences through the cytosol (White et al., 2023). In HeLa cells and yeast, the scale of heterogeneity in an ATP-related fluorescent signal within cells appears to be of a lower magnitude, though still present (Imamura et al., 2009; Takaine et al., 2019). The same probe was used to identify “hotspots” of ATP colocalized with sites of viral replication, with a 5-fold difference in ATP concentration between these hotspots and the cellular background (Ando et al., 2012) (Fig. 1C).

Our research question here can be thought of as the dual to these results, and an attempt to unite them. The above modelling and experimental work has characterized the existence and magnitude of cytosolic ATP gradients in *specific* cellular circumstances. Here we attempt to explore under what *general* cellular circumstances ATP gradients of a given magnitude could exist. To this end we require a system where many different instances can be explored in a computationally reasonable time. We therefore adopt a coarse-grained paradigm, considering a simplified 2D system that we design to retain the important quantitative characteristics determining the magnitude of ATP gradients while also being computationally tractable. For more detailed specific cellular circumstances, more precise modelling approaches may be desirable (Francis et al., 2024). We adopt the paradigm of “back-of-the envelope biology” (Johnston et al., 2014; Phillips & Milo, 2009), working with orders-of-magnitude quantities and physical reasoning to gain general numerical estimates of mechanism-linked quantities.

## Methods

### Equations of motion

Our basic picture is a reaction-diffusion model in a 2D model cell – that is, a model allowing ATP to diffuse and undergo reactions of production and consumption. Our governing partial differential equation is

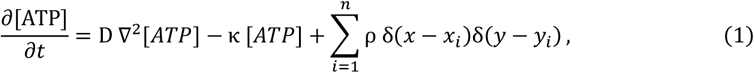

encoding the diffusion of ATP through the cell with diffusion constant D, the consumption of ATP with rate constant κthroughout the cell, and the production of ATP at rate ρ at a collection of discrete sites (*x*_*i*_,*y*_*i*_) corresponding to the *n* mitochondria in the cell. The consumption term may be applied uniformly throughout the cell or limited to a central region, as described in the text. We consider [ATP] to vary only in 2D, but picture a given cell depth to facilitate comparisons of concentration and volume; the third dimension is assumed to be relatively small and to have a uniform ATP profile (Fig. 1A-B). By default, we use a simple model cell of 50 μm × 50 μm × 10 μm (hence volume 2.5 × 10^4^ μm^3^) to model a (relatively large) plant cell (Fig. 1A-B) (Chustecki et al., 2021); we also consider 20 μm × 20 μm × 10 μm (hence volume 4 × 10^3^ μm^3^) to model the (relatively small) animal cell case (Puck et al., 1956), Bionumber 105879 (Milo et al., 2010). We also vary cell thickness to assess its contribution to ATP gradient magnitude.

It will immediately be seen that metabolism is effectively ignored in our model – we do not consider the details of the precursors to ATP production or of its consumption, and do not consider ADP, the concentration of which determines the free energy associated with the ATP landscape. For simplicity and biophysical generality, we just consider ATP as an independent diffusable metabolite with sources and sinks throughout the cell.

We also neglect ATP production from glycolysis. In conditions supporting oxidative phosphorylation, the rate of ATP production from mitochondria is over an order of magnitude higher (Mookerjee et al., 2017). Additionally, glycolytic ATP production will be more evenly spaced through the cell than mitochondrial ATP production, and hence contribute less to the establishment of spatial gradients of ATP concentration.

### Mitochondrial motion

In the case of motile mitochondria, we used two models for motion. Both models describe the physical change in a mitochondrial position vector **x**= (*x*_*i*_,*y*_*i*_) in one discrete timestep in our numerical simulation. First, random diffusion with diffusion constant scaled by ATP:

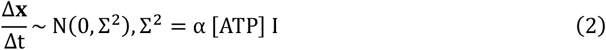

where *I* is the identity matrix and αis a scaling factor determining the scale of mitochondrial movement. This picture models ATP-dependent mitochondrial motion without systematic directionality. Second, directed motion down the [ATP] gradient:

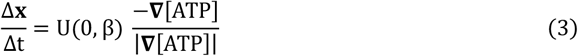

where βis a scaling factor determining the scale of mitochondrial movement, and the uniform random distribution is used both to model fluctuations in dynamics and for numerical convenience. The scaling factors we use are α= 5 × 10^-6^ and β= 1, chosen to restrict mitochondrial motion under typical ATP conditions to a maximum speed of order 1 μm s^-1^ (Chustecki et al., 2021). We ignore the locally-incurred ATP cost of moving mitochondria, which is assumed to be small compared to the other processes consuming ATP in the cell.

### Parameterisation

We use some parameter values (D) directly from previous observations. Others (κ,ρ)are less directly observable and are constrained through their indirect effect on observable quantities. For these, we assume some bounds on the orders of magnitude involved, and scan through values so that observed quantities take biologically reasonable values.

We assume cellular production of ATP is on the scale of 10^9^ ATP/s (Flamholz et al., 2014). We assume a default of 100 individual mitochondria in the cell, broadly compatible with observations in plants (Chustecki et al., 2021; Chustecki & Johnston, 2024; Logan, 2010; Logan & Leaver, 2000) (Fig. 1A) – counts vary dramatically across other species and tissue types, and mitochondria in other clades often fuse into networks (Aryaman, Johnston, et al., 2019; Hoitzing et al., 2015) (see Discussion). We vary this value to explore its influence on ATP gradient magnitude. We ignore glycolysis and other sources of ATP production, so that we expect a balancing ATP production per mitochondrion to be on the scale of 10^7^ ATP/s/mito. We impose an ATP diffusion rate of 2.5 × 10^2^ μm^2^ s^-1^ (Hubley et al., 1995); while muscle, the cell type for that observation, likely poses more restrictions to ATP diffusion than other cell types, diffusion in water is the same order of magnitude (7.1 × 10^2^ μm^2^ s^-1^ (Bowen & Martin, 1964)), so the range for this value appears reasonable.

Typical intracellular ATP concentrations are around 1-10 mM (Greiner & Glonek, 2021). A value of 1 mM corresponds to 10^-3^ mol dm^-3^ = 10^-3^ mol (10^15^ μm^3^)^-1^ = 10^-3^ × 6 × 10^23^ / 10^15^ = 6 × 10^5^ ATP / μm^3^. For our example model cell with a depth of 10 μm this corresponds to a ATP amount per square-micron region of around 6 × 10^4^ ATP / μm^2^.

### Numerics

We use a simple finite difference solver with a timestep of 2.5 × 10^-4^ s and grid element size of 0.5 μm, initializing the cell with zero ATP concentration throughout, and simulate the evolution of the system until a steady-state criterion is met (no relative ATP change greater than 10^-4^ across the domain). For comparison with biological criteria, we report the total consumption rate of ATP throughout the cell, and the total ATP amount in the cell.

As mentioned above, we work in a quasi-2D picture, where the depth of the cytoplasm is less important than the length and width. This is not unreasonable in, for example, plant hypocotyl cells, where the cytoplasm can be almost arbitrarily thin (Fig. 1A) (Chustecki et al., 2021; Lee Erickson et al., 2023). It can also be pictured as an approximation to cells which are relatively flat compared to their diameter, and where smaller ATP variability exists in the direction of depth than in other directions (Fig. 1B). In this situation, we neglect ATP variation in the depth direction and consider the 2D case where the concentration reflects an integrated value across cell depth.

## Results

### Inffuence of model source-sink distributions with static mitochondria

We first scanned through parameters (κ,ρ)(ATP consumption and production) to identify parameterisations compatible with biological values of cellular ATP content and consumption. We focus on parameterisations that, under equilibrium conditions, give cellular consumption values between 10^8^ ATP/s and 10^10^ ATP/s, and ATP concentration around 0.5-10mM (see Methods and (Flamholz et al., 2014; Greiner & Glonek, 2021)).

We consider several different arrangements of ATP sources (mitochondria) and sinks. We either randomly space mitochondria uniformly throughout the cell or cluster them in the centre; and we either space ATP consumption uniformly throughout the cell or restrict it to the central region (Fig. 2). In all cases, we run the simulated cell until ATP concentration reaches a steady state, then report the coefficient of variation (CV) – the ratio of the standard deviation of ATP concentration to the mean across in the cell (Fig. 3). Equilibration times were generally rapid under all parameterisations we considered. From an unphysical initial condition of zero ATP concentration, simulations typically reached a steady state ATP profile in under 30s of simulation time.

**Figure 2.**
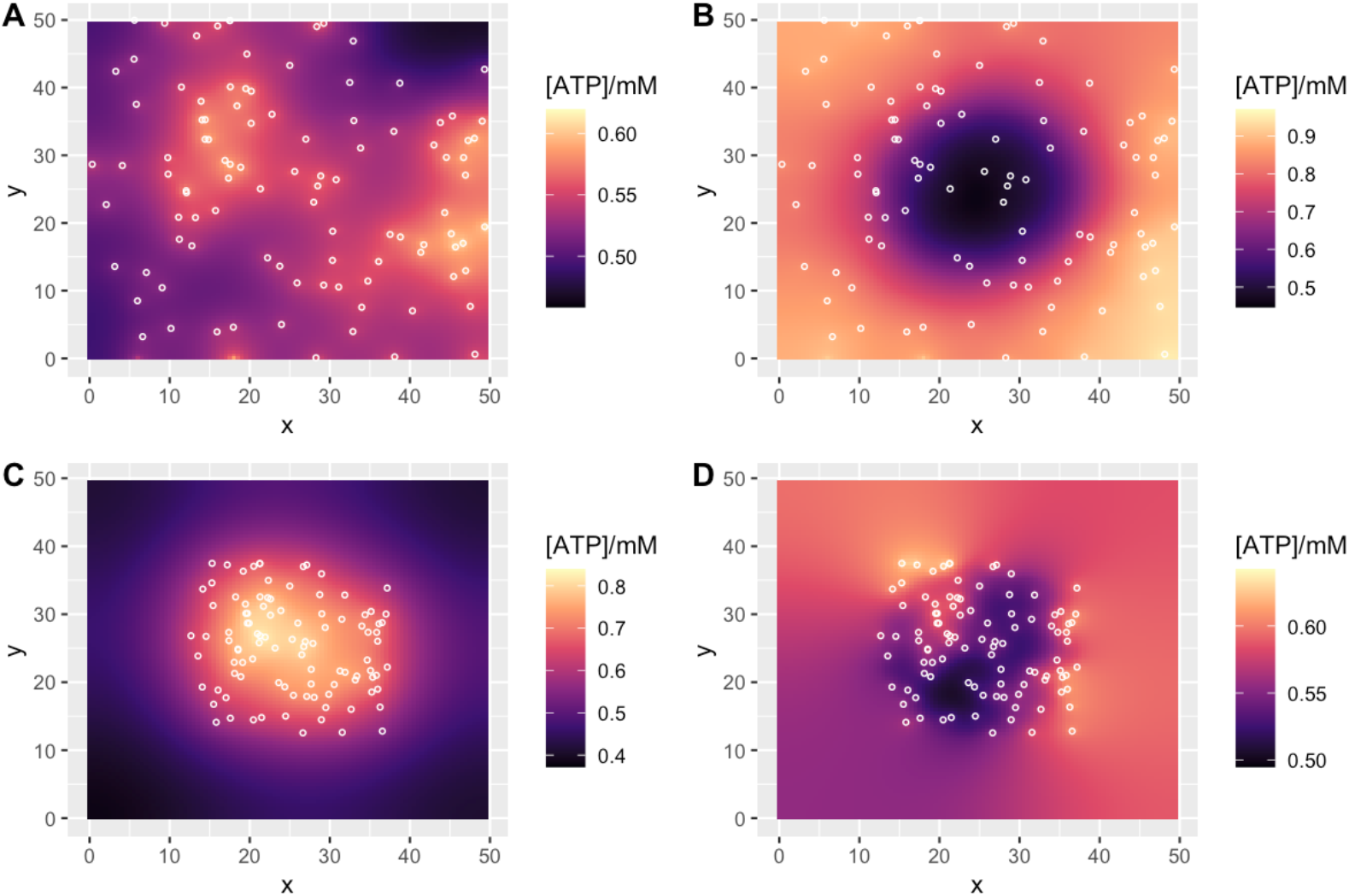
Examples of model behaviour with static mitochondria exhibiting ATP gradients. Long-term behaviour of ATP concentration in the cell under different cellular scenarios with static mitochondria. Plots give x-y dimensions of the model cell in μm, with local ATP concentration (heatmap) and positions of mitochondria (circles). (A) Uniform mitochondria, uniform ATP consumption; (B) uniform mitochondria, clustered ATP consumption; (C) clustered mitochondria, uniform ATP consumption; (D) clustered mitochondria, clustered ATP consumption. Parameterisations were chosen to give comparable ATP concentrations around 0.5mM in each case; they correspond to cellular consumption rates around 5.1 × 10^9^ molecules s^-1^.

**Figure 3.**
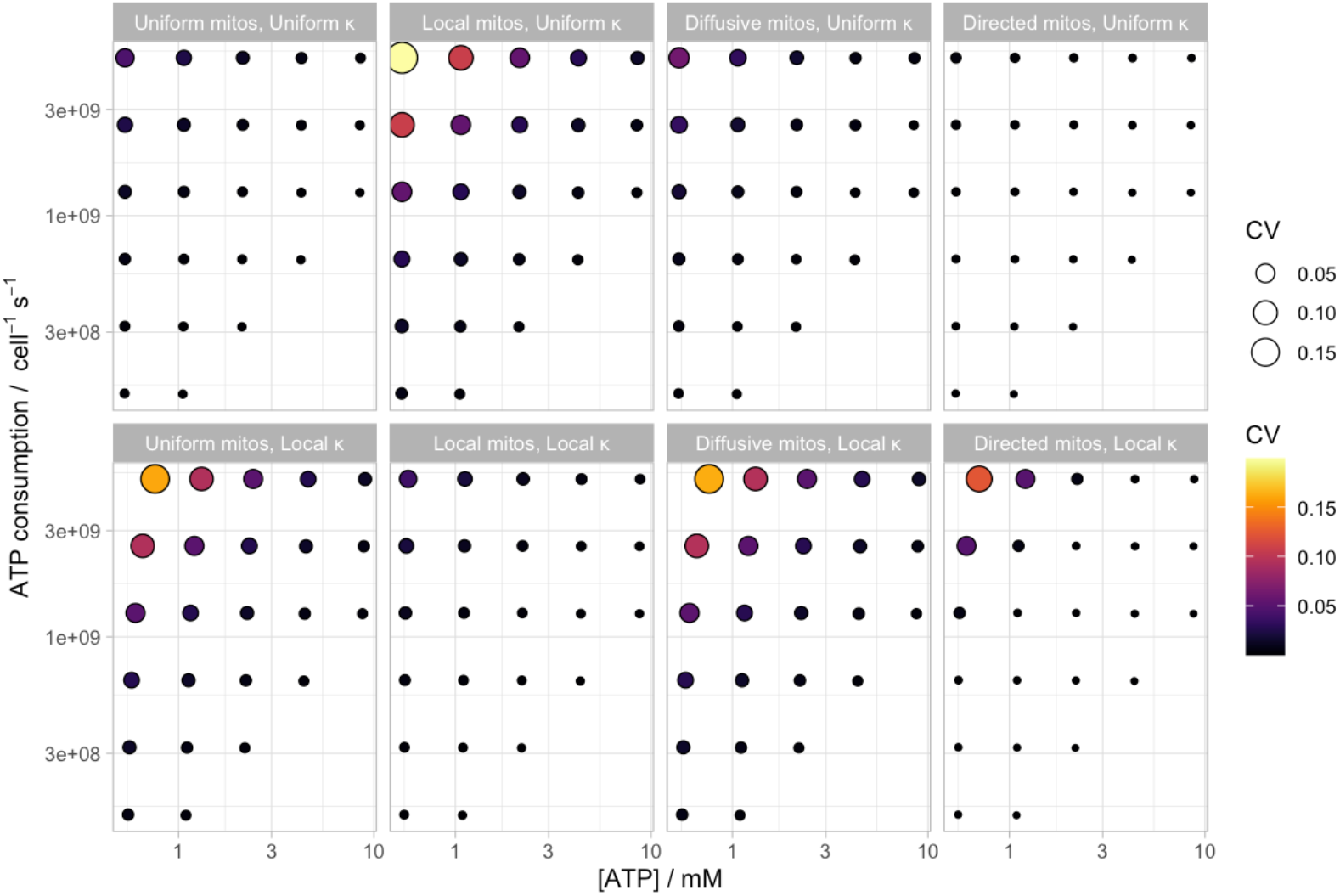
Model ATP concentration differences under different conditions. Coefficient of variation (CV) of ATP concentration across the cell with cellular ATP concentration and ATP consumption rate. Labels describe mitochondria (mitos) uniformly randomly distributed and static (uniform), clustered locally at the centre of the cell (local), undergoing diffusive motion (diffusive, Eqn. 2), and undergoing directed motion (directed, Eqn. 3); and ATP consumption (κ)uniform through the cell (uniform) or localized in the centre of the cell (local). These CV ranges correspond roughly to fold-change values between 0 and 2.5 (Supp. Fig. 1).

We found that strong determinants of ATP gradient magnitude were the overall concentration of ATP (lower concentrations supporting higher gradients) and the cellular rate of ATP consumption (higher consumptions supporting higher gradients). Substantial gradients with CVs over 10% typically occur only at the limits of our biological window for these values (Fig. 3). At given amounts of concentration and consumption, we found (following intuition) that the magnitude of ATP fold change dependent strongly on the arrangement of cellular sources and sinks. If static mitochondria and ATP consumption are “matched” – either evenly spaced or colocalised in a particular region through the cell, no combination of parameters compatible with biological observations led to more than a 10% CV in ATP across the cell, and most led to substantially smaller gradients (Fig. 2A, D; Fig. 3). However, if the positions of mitochondria and ATP consumption were less closely correlated, CV in ATP concentration across the cell could reach rather higher levels > 15% (Fig. 2B, C; Fig. 3). The CV was highly correlated with the fold range between the minimum and the maximum ATP concentrations found across the cell: these higher CV cases correspond to an over 2-fold difference between minimum and maximum ATP concentration (Fig. 2B, C; Supp. Fig. 1). These results did not change substantially when we considered a cell size more reflective of animal than plant cells (Supp. Fig. 2).

### Model behaviour with dynamic mitochondria

We next allowed mitochondria to move in the cell, following one of two models: random diffusion scaled by local ATP concentration (Eqn. 2), or directed motion down the local ATP gradient (Eqn. 3). Example dynamics and ATP profiles are shown in Fig. 4. No dynamic protocol qualitatively changed our results on ATP gradients (Fig. 3), although several emergent dynamics were clear. When mitochondria were actively moved against an ATP gradient, even spacing throughout the cell emerged in the case of uniform ATP consumption (Fig. 4C), following the observation of (Kajita et al., 2024) that the coupled production-motion system naturally leads to mitochondrial spacing. In the case of localised ATP consumption, mitochondria in this model moved preferentially towards the centre of the cell to set up “perinuclear” clustering with even spacing (Fig. 4D). Again, results were consistent in the case of smaller cells reflecting the more animal-like case (Supp. Fig. 2).

**Figure 4.**
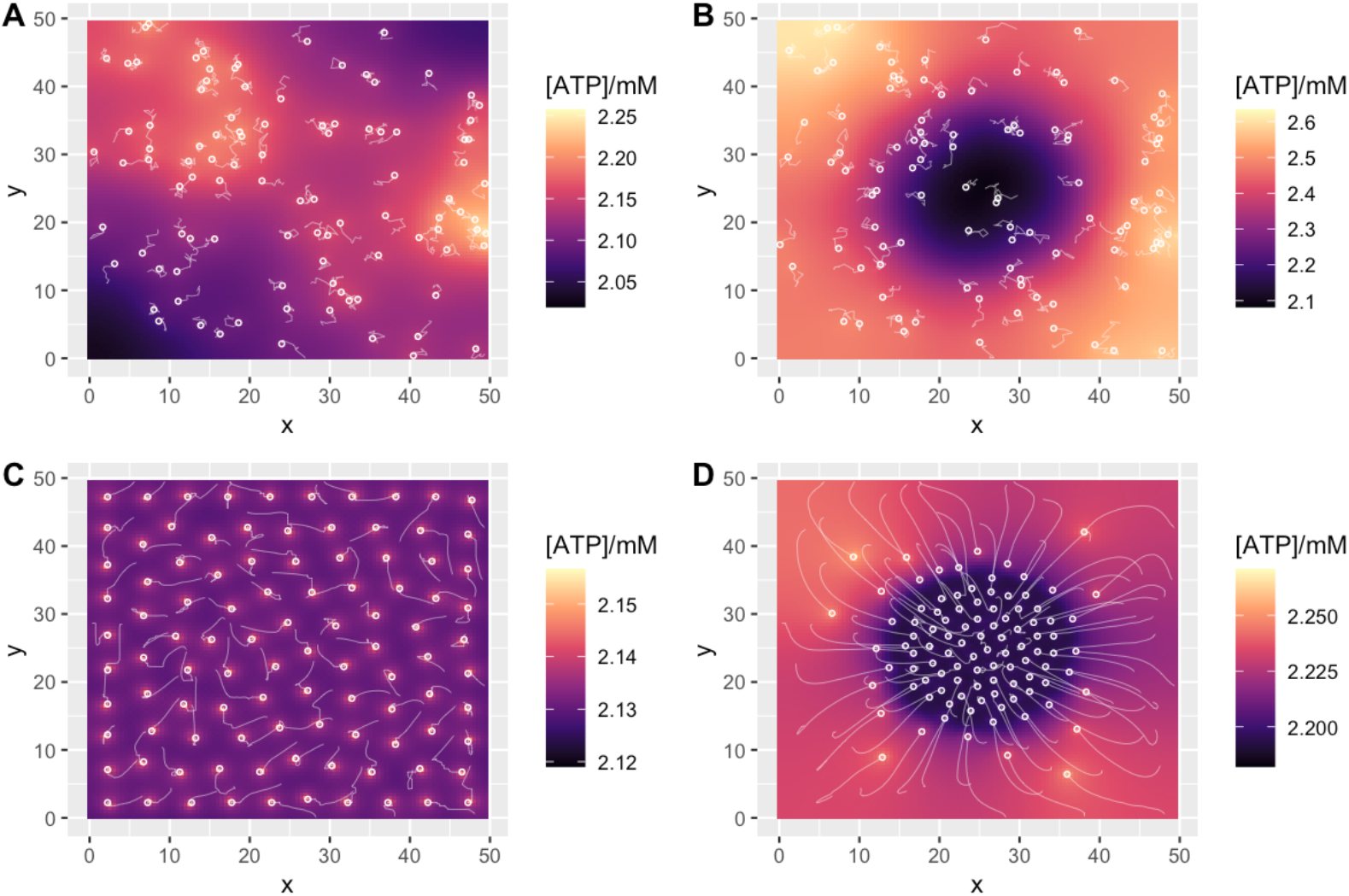
Model ATP behaviour with mitochondrial dynamics. Example long-term behaviour of ATP concentration in the cell under different cellular scenarios with dynamic mitochondria. Plots give x-y dimensions of the model cell in μm, with local ATP concentration (heatmap) and positions of mitochondria (circles). Light trails show mitochondrial motion. (A) Diffusive mitochondria, uniform ATP consumption; (B) diffusive mitochondria, clustered ATP consumption; (C) directed mitochondria, uniform ATP consumption; (D) directed mitochondria, clustered ATP consumption. For (A-B), trails show mitochondria motion over the final 10s of simulation; for (C-D), trials show mitochondrial motion throughout the simulation (from random initial conditions to a stable arrangement). Parameterisations were chosen to give comparable ATP concentrations around 2mM in each case; they correspond to cellular consumption rates around 5.1 × 10^9^ molecules s^-1^.

### Inffuence of model cell dimensions and mitochondrial number

We next asked how cell thickness and mitochondrial number influenced the behaviour of ATP gradients in the cell. To this end, we simulated model behaviour with fewer (50, Supp. Fig. 3) and more (200, Supp. Fig. 4) mitochondria, and with thinner (2 μm, Supp. Fig. 5) and thicker (20 μm, Supp. Fig. 6) dimensions in the “depth” direction. In each case the qualitative direction of the results is the same – lower ATP concentrations and higher consumption rates give higher CVs for ATP, and localised energy demand without compensatory mitochondrial localisation gives the highest CV magnitudes.

The number of mitochondria in our model did not lead to a notable change in the scale of model CVs at a given concentration-consumption state (Fig. 3, Supp. Figs. 3A, 4A). For example, the CV for diffusive mitochondria and centralised demand at an ATP concentration of 1mM and a consumption rate of 10^9^ molecules s^-1^ is typically around 0.05 for all the mitochondrial numbers we consider; the most extreme case shared by all experiments, with clustered mitochondria, uniform demand, [ATP] around 0.5mM and consumption 3 × 10^9^ s^-1^, had a CV around 0.12 in all cases. The thickness of the cell in our model had a stronger effect. At 1mM concentration and 10^9^ molecules s^-1^ consumption, the thinner cell had a CV around 0.25 and the thicker cell around 0.02, suggesting an inverse linear scaling of CV with cell thickness.

We verified that for a given (κ,ρ)parameterization of the model, the cell depth did not influence the number of ATP molecules per simulation element (as the cell depth does not feature in the simulation itself, only playing a role in the *post hoc* calculation of concentration from this quantity). However, a given number of ATP molecules per model area element will translate to a higher concentration in a thinner cell and a lower concentration in a thicker cell. The (κ,ρ)parameterization corresponding to a given cellular ATP concentration will thus depend on cell thickness – broadly, thinner cells achieve a given ATP concentration with lower production or higher consumption terms. The increase in CV at lower model thicknesses for a given cellular ATP concentration can then be explained by the change in the rates of the production and consumption processes needed to achieve that concentration for a given cell thickness.

## Discussion

In this simple biophysical model, after the concentration and consumption of ATP, the relative positioning of ATP sources and sinks strongly influences whether substantial ATP concentration gradients exist in the cell. Under reasonable biological parameterisations, it is possible either for negligible differences (CV < 1%) to exist across the cell, or to have fairly substantial concentration differences (CV > 15%, 2-fold changes across the cell). A notable example of this structure-induced difference is in the low concentration, high consumption region of Fig. 3 – with static, clustered mitochondria and uniform consumption, CV exceeds 15% (as in Fig. 3C), while with dynamic mitochondria that respond to the ATP gradient, CV is almost negligible (resembling Fig. 4C). The specific scales of CV depend on the thickness of the model cell, with thinner cytoplasmic sections supporting proportionally higher CV values, and vice versa. The dependence of CV on mitochondrial number (and density) is much more limited. Across all models, CVs are highest when ATP concentrations are lowest and consumption rates are highest. Agreeing with experimental observations (Matsumoto et al., 2022; Schuler et al., 2017) and intuition, the highest ATP concentrations are found in regions of densely arrangement mitochondria.

Heterogeneity in ATP profile (where it existed) had a substantial effect on the motion of mitochondria. In the case of directed motion down the ATP gradient, mitochondria rapidly adopted positioning that reflected the spread or localisation of demand (Fig. 4C-D). We saw little evidence under passive diffusion that differences in ATP concentration across the cell led to pronounced emergence of mitochondrial structure – for example, there was little evidence that slower mitochondrial motion in lower-concentration regions of the cell led to a “buildup” of mitochondria in those regions (Fig. 4B). Active transport of mitochondria down the concentration gradient was required for such emergent structure. In this situation, we particularly saw the emergence of even spacing regardless of the geometry of the region of demand – matching the “active thermodynamic force” driving even spacing from (Kajita et al., 2024). This spatial heterogeneity in macroscopic cellular components, emerging from variability in an underlying concentration field, underlines the potential importance of considering spatial profiles in “whole-cell modelling” pictures (Goldberg et al., 2018). Work considering the optimal arrangement of enzymes using principles of optimisation (and operations research) suggests a promising framework for such consideration (Chustecki & Johnston, 2024; Giunta et al., 2022; Sutherland, 2005).

Our model is obviously a very coarse-grained representation of a cell. ATP metabolism is represented simply by point production and proportional degradation, with no representation of ADP, mitochondrial heterogeneity (Aryaman, Johnston, et al., 2019), or other degrees of freedom. The geometry of the cell is fixed and the third dimension is effectively ignored.

However, we believe this simplified picture both captures the most essential drivers of ATP concentration gradients and retains interpretability without including the large number of additional parameters that would be required to capture these effects. Modelling approaches at a finer-grained microscopic scale are being developed (Francis et al., 2024) and provide an alternative to our general mesoscopic picture for specific cellular circumstances.

In different specific situations, mitochondria will not be the dominant source of ATP in the cell. Green plant tissues in the light, for example, generate substantial ATP from chloroplasts (also localised organelles, though often tightly packed throughout the body of such cells), In low-oxygen conditions or nutrient substrates that do not support OXPHOS, other processes like glycolysis may be the more important sources of ATP.

The fragmented nature of our model mitochondria is, of course, not representative of all cell types in all organisms. While mitochondria exist in a predominantly fragmented form in, for example, somatic plant cells (Logan, 2010; Logan & Leaver, 2000), in other organisms they fuse into larger connected networks (Hoitzing et al., 2015). If mitochondria instead form a reticulated network along which ATP production is relatively uniform, ATP sources will be more evenly spread through the cell and concentration gradients will be diminished further. Generalisation of this simple model to support more connected mitochondrial elements and/or elements beyond simple point sources could readily address this connected picture in future.

## Supplementary Information

**Supplementary Figure 1.**
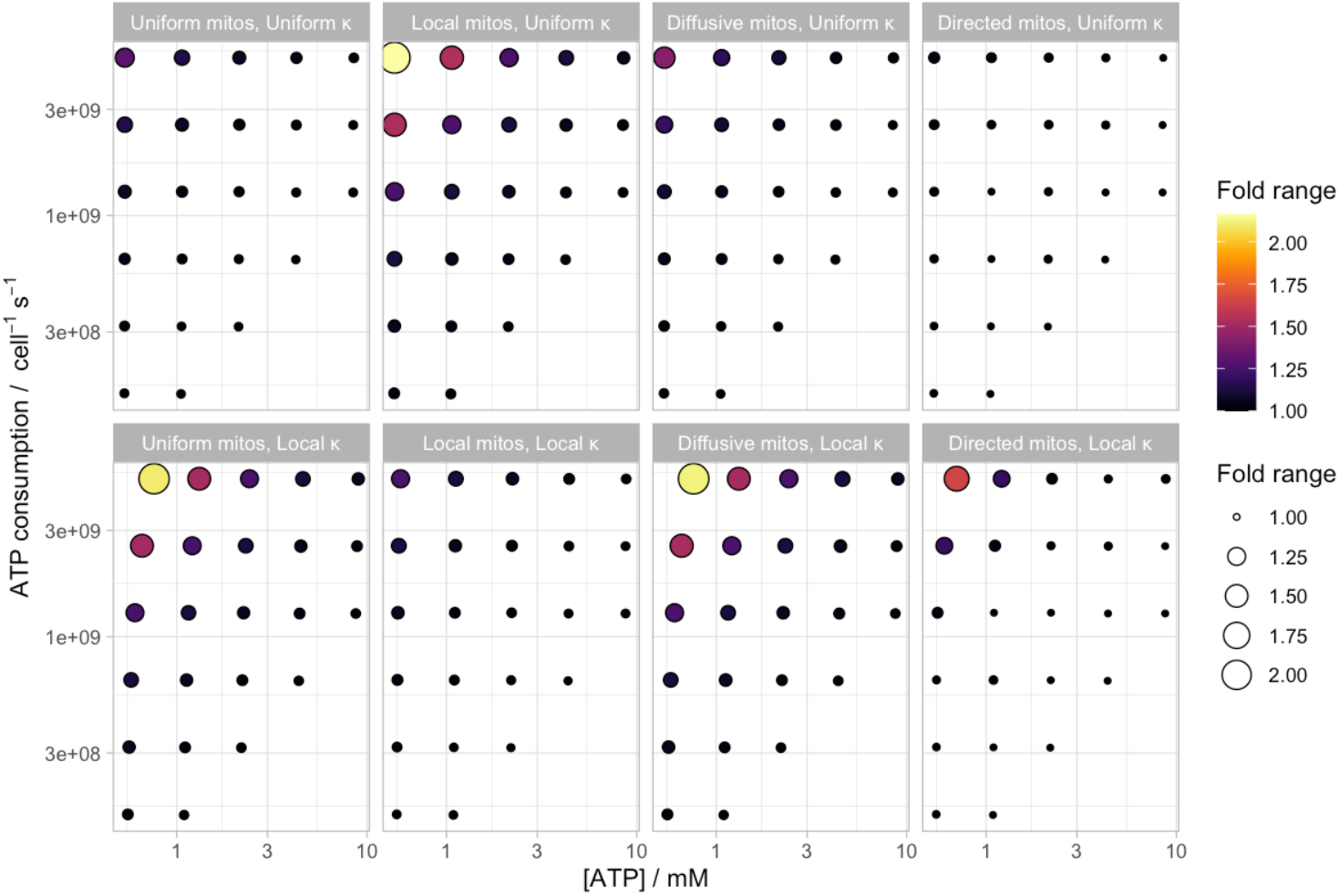
Fold-change in ATP concentration. ATP concentration gradients plotted as fold change between minimum and maximum values in the cell, rather than CV as in the main text.

**Supplementary Figure 2.**
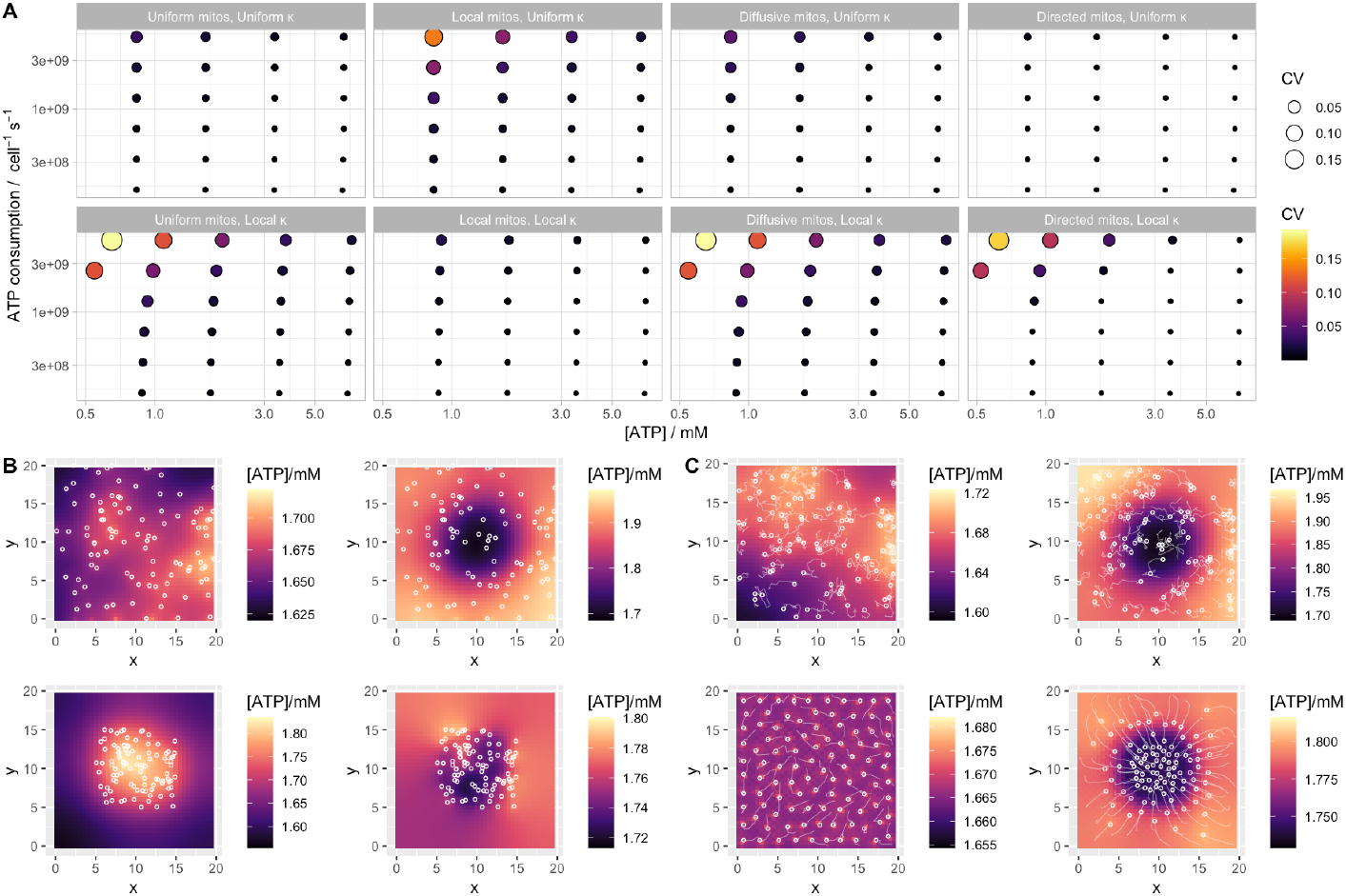
Animal cell version of the model. This approach uses a model cell 20 μm × 20 μm × 10 μm (hence volume 4 × 10^3^ μm^3^). (A) CV of ATP concentration as in Fig. 3. (B) Long-term concentration profile with static mitochondria as in Fig. 2. (C) Long-term concentration profile with static mitochondria as in Fig. 4. Parameters in (B-C) correspond to cellular consumption rates around 2.6 × 10^9^ molecules s^-1^.

**Supplementary Figure 3.**
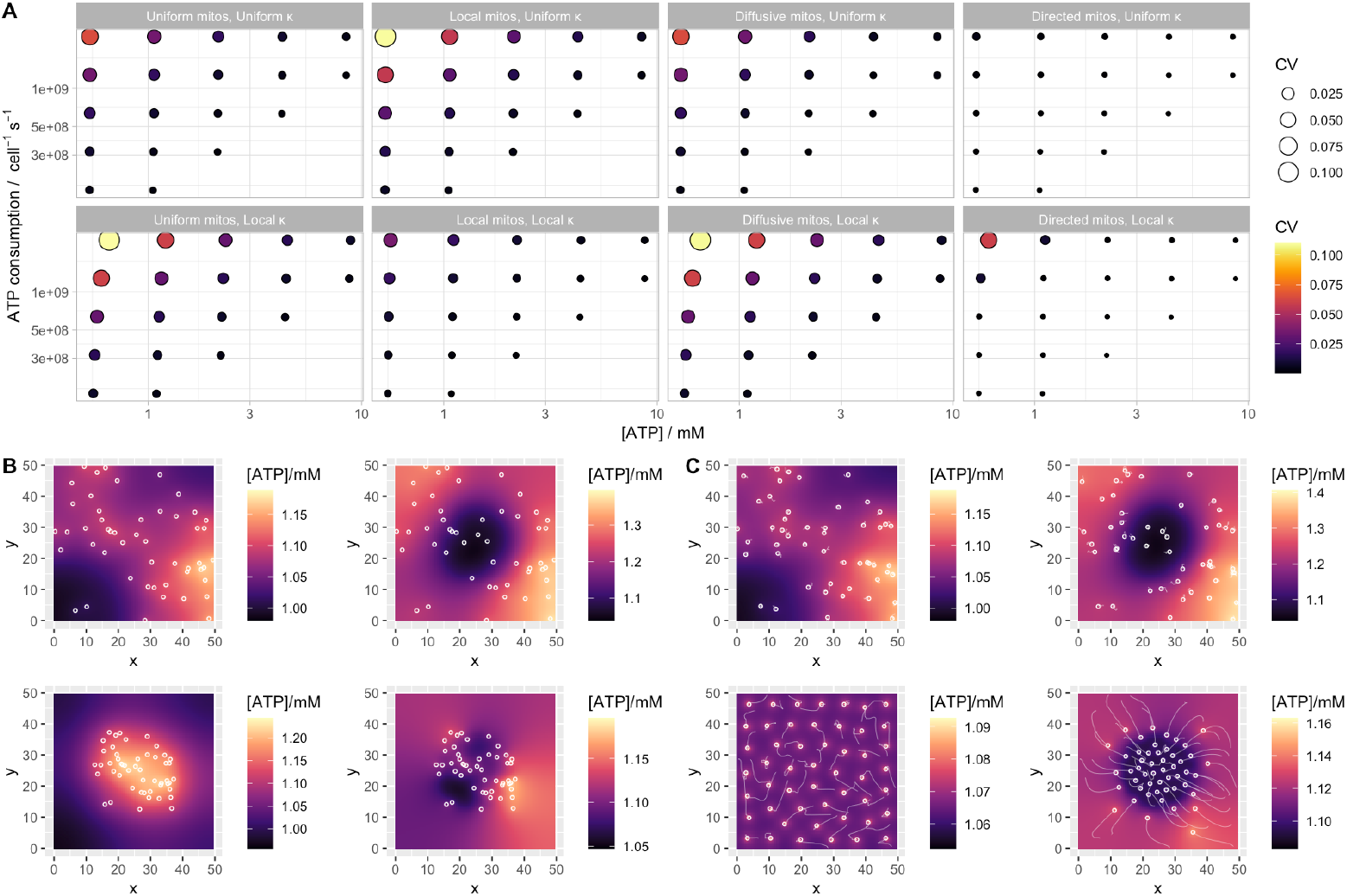
Model with fewer mitochondria. 50 mitochondria in the simulated cell. (A) CV of ATP concentration as in Fig. 3. (B) Long-term concentration profile with static mitochondria as in Fig. 2. (C) Long-term concentration profile with static mitochondria as in Fig. 4.

**Supplementary Figure 4.**
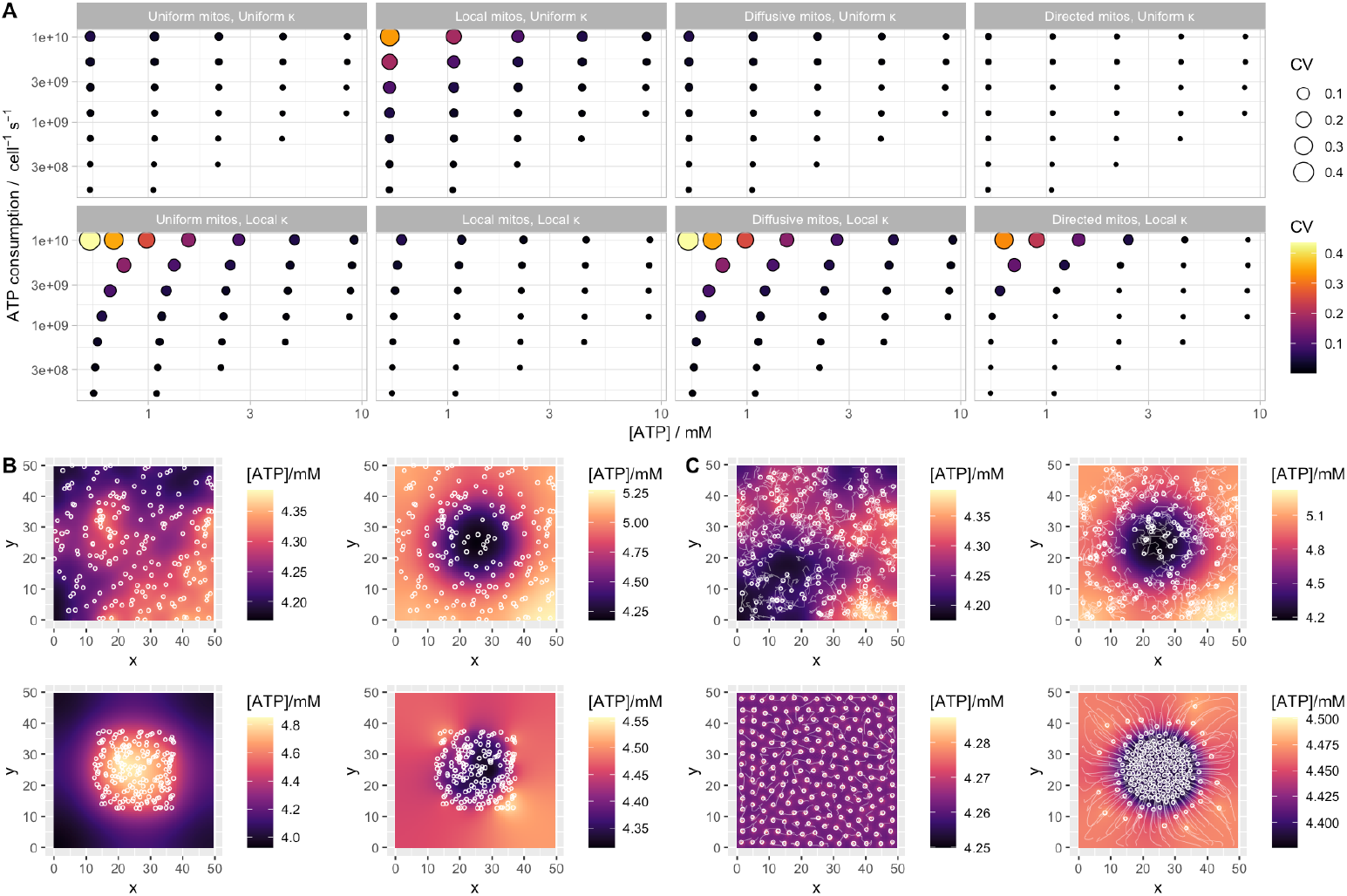
Model with more mitochondria. 200 mitochondria in the simulated cell. (A) CV of ATP concentration as in Fig. 3. (B) Long-term concentration profile with static mitochondria as in Fig. 2. (C) Long-term concentration profile with static mitochondria as in Fig. 4.

**Supplementary Figure 5.**
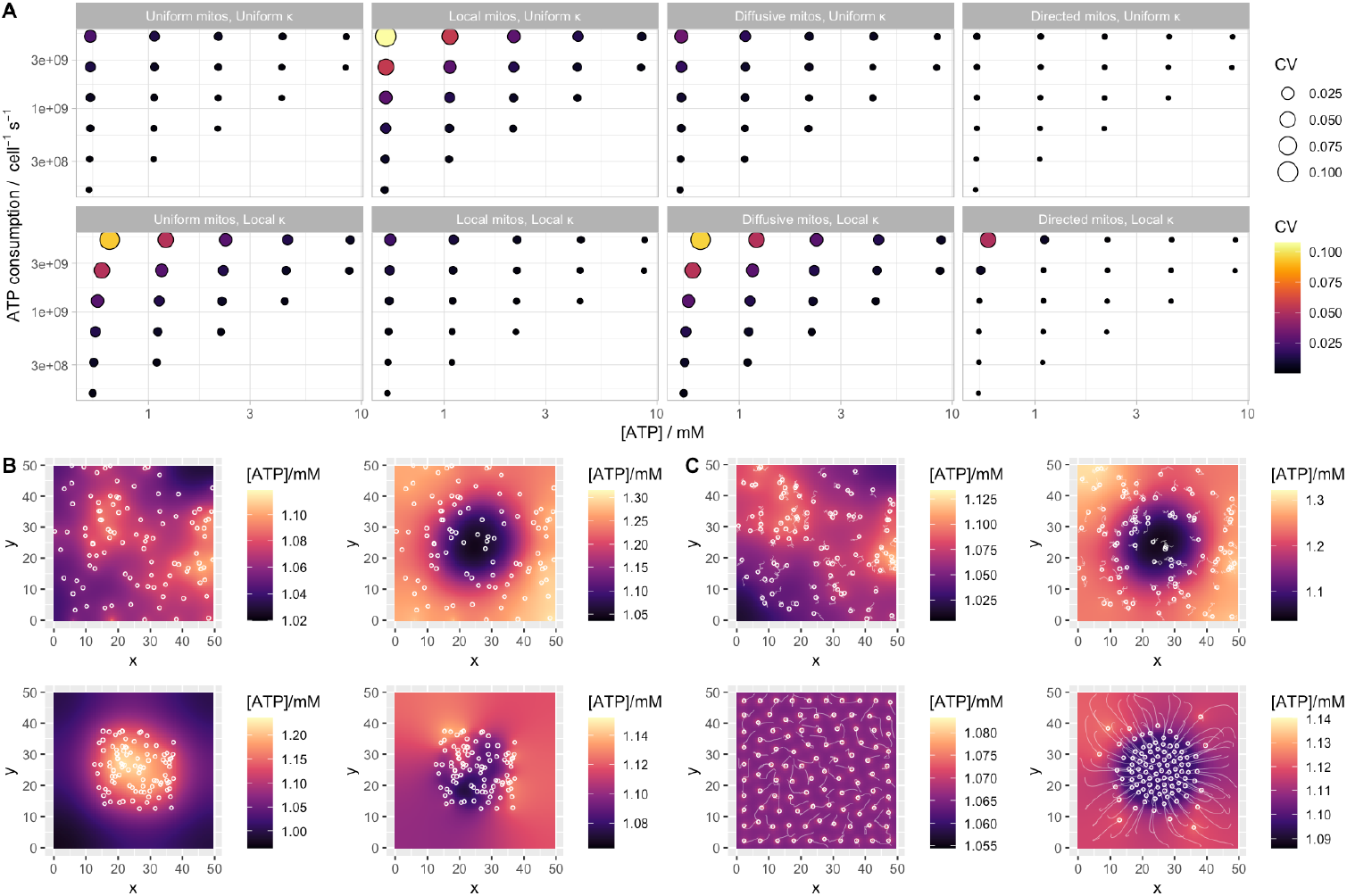
Model with higher cell thickness. Cell thickness assumed to be 20μm. (A) CV of ATP concentration as in Fig. 3. (B) Long-term concentration profile with static mitochondria as in Fig. 2. (C) Long-term concentration profile with static mitochondria as in Fig. 4.

**Supplementary Figure 6.**
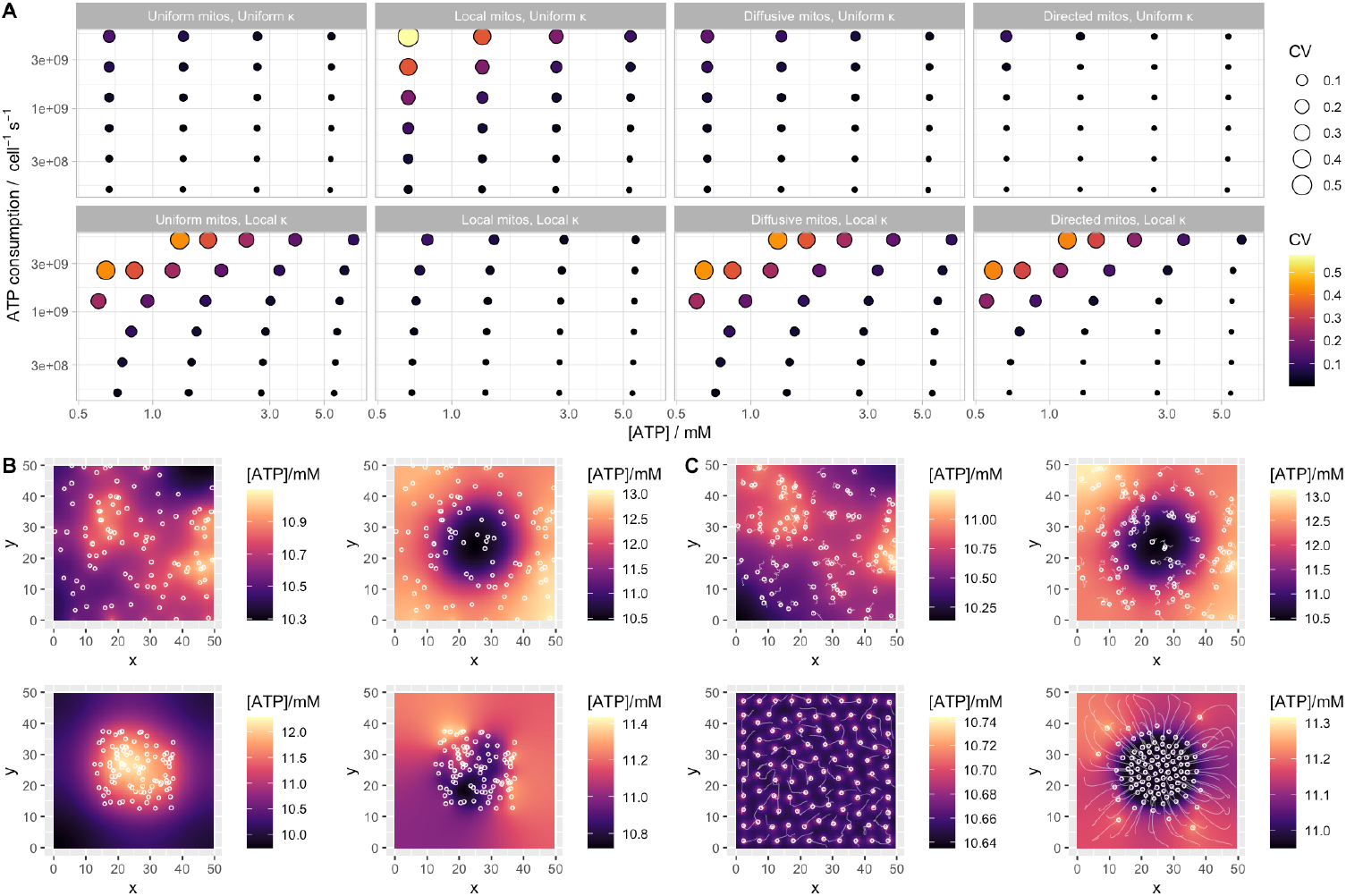
Model with lower cell thickness. Cell thickness assumed to be 2μm. (A) CV of ATP concentration as in Fig. 3. (B) Long-term concentration profile with static mitochondria as in Fig. 2. (C) Long-term concentration profile with static mitochondria as in Fig. 4.

